# Bacteriophage-mediated decolonization of *Enterobacteriaceae* in a novel *Galleria mellonella* gut colonization model

**DOI:** 10.1101/2023.03.27.534343

**Authors:** Kamran A. Mirza, Sebastian Jacob, Sandor Nietzsche, Oliwia Makarewicz, Mathias W. Pletz, Lara Thieme

**Affiliations:** Jena University Hospital, Institute of Infectious Diseases and Infection Control, Friedrich-Schiller-University Jena, Am Klinikum 1, 07747 Jena, Germany; Jena University Hospital, Leibniz Center for Photonics in Infection Research, Friedrich Schiller University Jena, 07747 Jena, Germany; Electron Microscope Center, Jena University Hospital, Ziegelmühlenweg 1, 07743 Jena, Germany

**Keywords:** Larvae, Force-feed, Phage treatment, 3Rs, Invertebrate model

## Abstract

**Purpose:** *Galleria mellonella* larvae have emerged as an invertebrate model for studying bacterial pathogenesis and novel therapeutic options due to ethical concerns associated with the use of mammalian models such as mice. The benefits of using *G. mellonella* larvae include a less complex microbiome in the gut, making it suitable for gut colonization studies. The intestinal colonization of *Klebsiella pneumoniae* (Kp) and *Escherichia coli* (Ec), two of the most antibiotic-resistant bacteria on the World Health Organization’s (WHO) priority list, plays a key role in the spread of antibiotic resistance. Bacteriophage therapy is emerging as a promising alternative for antibiotic-resistant bacteria due to its ability to specifically target and infect bacterial hosts, making it suitable for gut decontamination. This study aimed to establish a novel *Enterobacteriaceae G. mellonella* larvae gut colonization model and compare the efficacy of conventional antibiotic treatment with a one-time phage cocktail in decolonizing the gut.

**Approach:** Larvae were force-fed with different concentrations of bacterial doses of *K. pneumoniae* and *E. coli* at 0 h, 24 h, and 48 h, followed by survival monitoring at 24 h intervals. After 48 h and 120 h of the last force feed, the colony forming unit (CFU) count in the gut was evaluated. After successful colonization, larvae were one-time force-fed with either a 10^7^ PFU/larvae bacteriophage cocktail or with ciprofloxacin 4 mg/L or meropenem 2 mg/L. After 24 h post phage feeding, CFU counts were determined.

**Main findings:** Three bacterial doses of 10^6^ CFU/larvae led to a stable gut colonization in the larvae gut regardless of the *K. pneumoniae* and *E. coli* strains. Bacteriophage force-feeding reduced bacterial colonization by 4 log_10_ CFU/larvae whereas antibiotic treatment led to a 2 log_10_ CFU/larvae reduction compared to the control. The novel alternative *G. mellonella* model for gut colonization studies can be used for proof-of-concept studies, reducing or even obviating the number of follow-up experiments in vertebrate models.

## 1 Introduction

The intestine is a key reservoir of highly antibiotic-resistant, opportunistic pathogens such as bacteria of the *Enterobacteriaceae* family. *Klebsiella pneumoniae* and *Escherichia coli* have been identified by the World Health Organization (WHO) as high-priority pathogens within the *Enterobacteriaceae* family due to their resistance to the last resort antibiotics, the carbapenems, leading to high mortality in patients and high risk of hospital outbreaks (Tacconelli et al., 2018). Intestinal colonization with carbapenemase producing *Enterobacteriaceae* (CPE), such as *K. pneumoniae* and *E. coli*, is associated with an increased risk of subsequent nosocomial infections, e.g., pneumonia, urinary tract infection or sepsis. Patients with long-term intensive care unit hospitalization are at high risk of CPE intestinal colonization and breakthrough infections due to weak mucosal immunity, advanced age and prolonged uptake of antibiotics (Halpin et al., 2016). Thus, selective mitigation of the gut from CPE might prevent downstream lethal infections. However, many CPE are resistant to other antibiotic classes suitable for gut treatment. Further, antibiotics lack host specificity resulting in dysbiosis of the gut microbiome and increased risk of difficult-to-treat infections by *Clostridium difficile* (Ghose, 2013). An alternative approach is therefore the therapeutic application of bacteriophages.

Bacteriophages are bacterial viruses that effectively kill bacteria during their lytic cycle. They can have very high specificities, not only for one bacterial species, but also for a certain subspecies or even strain, and thus represent a highly interesting type of treatment to specifically address a certain pathogen (Gordillo Altamirano & Barr, 2019). Although bacteriophages were discovered over a century ago, the last decade has played a crucial role in exploring their therapeutic potential. Bacteriophages possess several benefits over antibiotics, such as host specificity, which means it does not affect the microbiota or mammalian cells (Gordillo Altamirano & Barr, 2019). Another benefit is self-limitation, which is the cessation of bacteriophage activity when the bacterial host is eliminated (Gordillo Altamirano & Barr, 2019). Bacteriophages were successfully applied intranasally and topically in mice against *K. pneumoniae*-associated lung and wound infections, revealing decreased bacterial loads and mortality rates (Cao et al., 2015) (Kumari et al., 2011). A recent study also identified two lytic phages capable of intestinal decolonization of *K. pneumoniae* in a mouse model (Fang et al., 2022).

To overcome strong ethical concerns due to high burden of disease in mammal infection models, researchers have reached alternative invertebrate models, such as the *Galleria mellonella* larvae infection model. Following the 3Rs framework in animal research (Replacement, Reduction, Refinement), larval models allow for high throughput and proof-of-concept screens, leading to a reduction of the number of animals in subsequent mammalian experiments. Larval research models in general are associated with low costs and are easy to handle. Thus, in contrast to mammalian experiments, no specialized training, complex animal husbandry and ethical approval are required. The particular advantages of *G. mellonella* larvae compared to other invertebrate models include a comparatively similar innate immune response to mammals, susceptibility to human pathogens, and survival at a temperature range of 37 to 42 °C. *G. mellonella* larvae can be easily inoculated orally or *via* their abdominal proleg structure. Their gradual melanization as an immune response to a pathogen allows easy monitoring of the course of infection in virulence studies and in efficacy studies of antimicrobial agents.

Many studies have used the *G. mellonella* model to evaluate antimicrobial efficacies against various pathogens in larval hemolymph, i.e., mimicking a bloodstream infection, including the efficacy of bacteriophages against *K. pneumoniae* strains (Gorodnichev et al., 2021; Insua et al., 2013; Sugeçti, 2021; Thiry et al., 2019). Studies of a selective gut decolonization from CPE by phages are missing due to the general lack of a well-standardized *G. mellonella* gut colonization model. Therefore, this study aimed to establish a stable gut colonization by *K. pneumoniae* and *E. coli* as most common CPEs in *G. mellonella* and to evaluate the effect of an oral bacteriophage treatment on the reduction of the CPE load in the gut.

## 2 Animals, materials and methods

### 2.1 Bacterial strains and bacteriophages

Two clinical isolates and one laboratory standard of each species were used for the studies (Table 1). The clinical isolates were previously collected in studies approved by the ethical committee of the Jena University Hospital (3852/07-13 and 3694-02/13). The minimum inhibitory concentrations (MICs) of the antibiotics were routinely assessed according to the by European Committee of Antimicrobial Susceptibility Testing (EUCAST) via VITEK2 (bioMérieux, Marcy-l’Étoile, France) and interpreted according to the clinical EUCAST breakpoints in 2022.

**Table 1.** Bacteria strains used for colonization studies and their characteristics.

|  | *K. pneumoniae* |  |  | *E. coli* |  |  |
| --- | --- | --- | --- | --- | --- | --- |
| Strain | ATCC 700603 | Kp1452 0 | Kp 41961 4 | ATCC 35218 | Ec 208873 | Ec 280624 |
| Specimen | LS | Urine | Stool | LS | Urine | Stool |
| Amoxicillin | 8 | 32 (R) | - | 8 | 32 (R) | 32 (R) |
| Fosfomycin | - | 64 (R) | - | - | 64 (R) | 64 (R) |
| Piperacilline/tazobactam | 16 | 16 (I) | 128 (R) | 1 | 4 (I) | 128 (R) |
| Ceftazidime | 32 | 32 (R) | 64 (R) | - | 4 (R) | 64 (R) |
| Meropenem | <0.25 | 0.25 (S) | 2 (R) | - | 0.25 (S) | 2 (R) |
| Ciprofloxacin | 0.5 | 4 (R) | 4 (R) | - | 4 (R) | 4 (R) |
| Gentamicin | 8 | - | 8 (R) | - | 1 (S) | 1 (R) |

LS = laboratory standard, sensitive (S), resistant (R), intermediate (I)

The bacteriophages UZG4 and UZG13 (Table 2) and their propagation host *K. pneumoniae* 1711-O4741 strain were obtained from the Ghent University Hospital, Belgium.

**Table 2.**
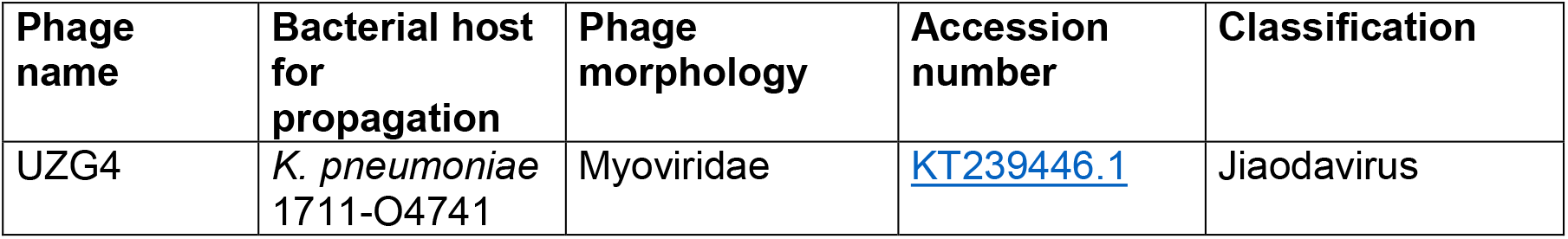
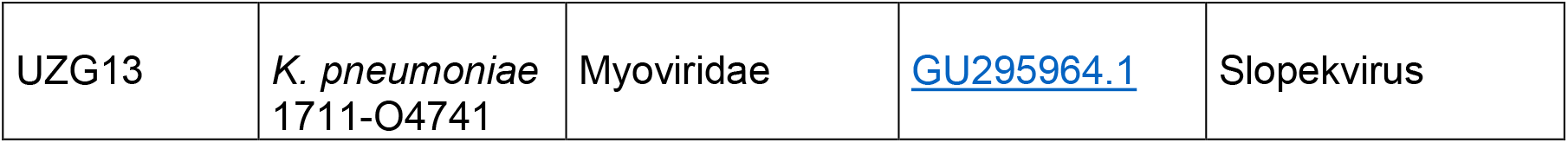
Bacteriophages used in the study.

| Phage name | Bacterial host for propagation | Phage morphology | Accession number | Classification |
| --- | --- | --- | --- | --- |
| UZG4 | *K. pneumoniae* 1711-O4741 | Myoviridae | KT239446.1 | Jiaodavirus |
| UZG13 | *K. pneumoniae*<br>1711-O4741 | Myoviridae | GU295964.1 | Slopekvirus |

### 2.2 Culturing and selection of bacteria

Bacteria were cultured in Müller Hinton (MH) broth (Oxoid Deutschland GmbH, Wiesel, Germany) at 37°C in a shaking incubator for 2 hours to reach the early exponential phase. Bacteria in the early exponential phase were used throughout the force-feeding experiments. MacConkey agar (Becton Dickinson, New Jersey, USA) was used as the selective media for the selection of Gram-negative *K. pneumoniae* and *E. coli* from the commensal microbiome of larvae gut, which is predominantly Gram-positive *Enterococcus* species (Allonsius et al., 2019).

Ciprofloxacin (PanReac AppliChem, Darmstadt, Germany) and meropenem (TCI Germany GmbH, Eschborn, Germany) were freshly prepared as 10 mg/mL stock solutions in sterile distilled water before use and for larvae treatment 4 mg/L ciprofloxacin and 2 mg/L were used in MH broth.

### 2.3 Phage titration

A culture of *K. pneumoniae* 1711-O4741 was grown overnight at 37 °C and 160 rotations per minute (RPM). In a 15 mL tube, 300 µL of bacterial suspension was mixed with 100 µL of the phage lysate and incubated at 37 °C and 160 RPM for 30 min. After the incubation, 4 mL of top agar (0.5 % agar in Luria-Bertani (LB) broth (Oxoid Germany GmbH)) was added to the mixture, vortexed and quickly and uniformly poured onto LB agar (Oxoid Germany GmbH) plate. The plate was incubated at 37 °C overnight. Subsequently, 5 mL of LB broth was pipetted onto the agar plate and incubated at 37 °C with 80 RPM in an orbital shaker for 2 h. After the incubation, the lysate and the top agar were harvested and centrifuged at 10000 relative centrifugal force (RCF) for 5 min. The supernatant was collected and filtered *via* a 0.2 µm syringe filter (Buch & Holm A/S, Herlev, Denmark). The phage titer was evaluated using double agar overlay assays (Kropinski et al., 2009). The phage stocks were adjusted in LB broth to a titer of 10^10^ plaque forming units (PFU)/mL and stored at 4°C.

### 2.4 Galleria mellonella

*G. mellonella* wax moth larvae were obtained from Bruno Mariani-FLOTEX (Augsburg, Germany). Larvae were stored in a refrigerator at 15 °C temperature for maximum of 2 weeks. Throughout the experiments, larvae of an average size of 3.3 cm (with a standard deviation (SD) of ±0.12 cm) were used.

### 2.5 Gut colonization of *G. mellonella* larvae

For the proof of concept, a bacterial suspension of Kp 14520 was taken from early exponential phase and adjusted to 10^8^ CFU/mL in MH broth. Larvae in group sizes of n = 20×3 per bacterial dose were fed with logarithmically increasing doses of bacteria from 10^2^ to 10^6^ CFU/larvae. Force-feeding was performed each at 0 h, 24 h, and 48 h using a 10 µl ALS syringe (Agilent Technologies, Santa Clara, USA) under a Stemi 2000-C microscope (Carl Zeiss, Oberkochen; Germany) (Figure S1) followed by incubation at 37°C (Figure 1). Survival of the larvae was monitored at 24 h, 48 h and 72 h post-force-feeding and Kaplan-Meier curves were generated. Post 24 h and 48 h of last force-feeding, i.e., 72 h and 96 h after the first dose of bacteria, larvae were anesthetized on ice for 30 min and the gut was surgically removed and transferred into 300 µL of 1 x phosphate buffer saline (PBS) (Carl Roth GmbH, Karlsruhe, Germany). The gut was crushed thoroughly using dissecting needle lancet, vortexed for 1 min, serially diluted, and plated as follows. For CFU counting, MacConkey agar plates were freshly prepared and used. To check the inter-strain variability in intestinal colonization, selected bacterial doses of two further *Klebsiella* strains, i.e. 10^5^ and 10^6^ CFU/larvae of Kp 419614 and ATCC 700603, were force-fed to the larvae following the same protocol. The larvae were also colonized with *E. coli* ATCC 35218, isolates Ec 280624 and Ec 208873 to check the variation among different *Enterobacteriaceae* species.

**Figure 1.**
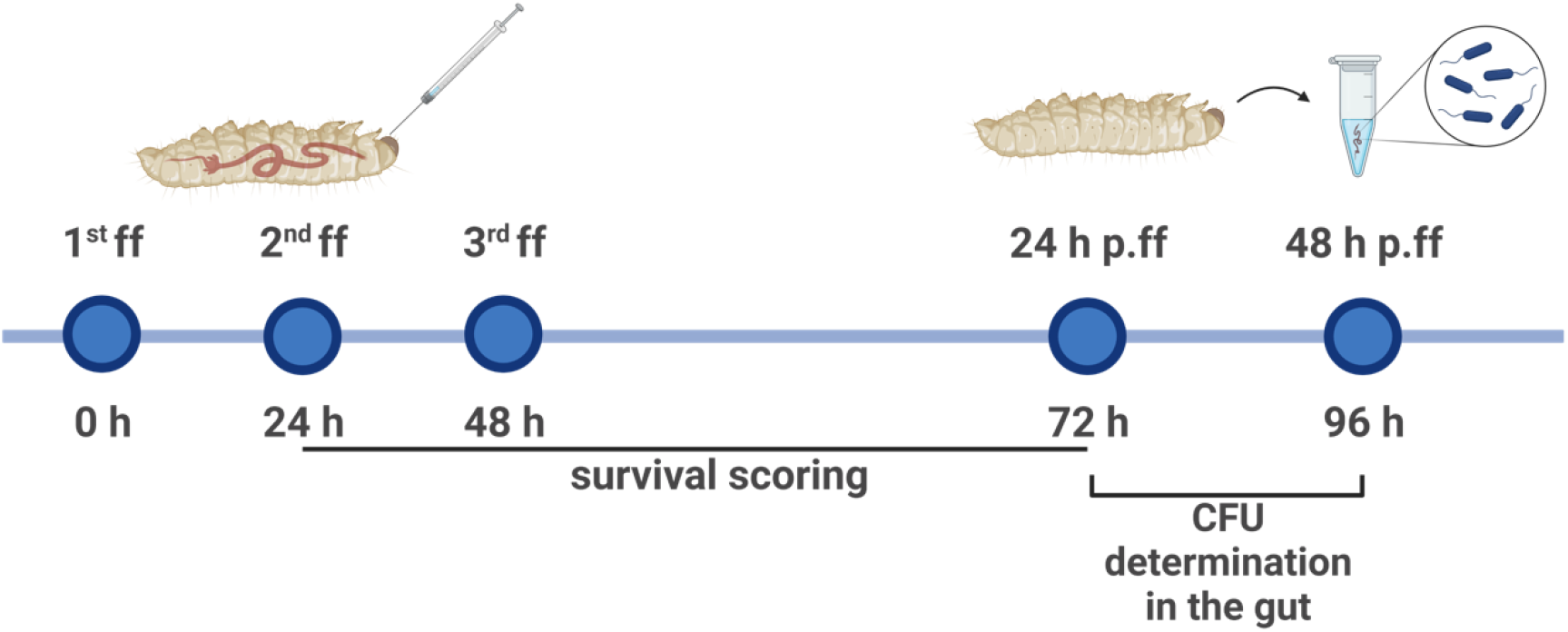
Timeline of the methodology adopted for the establishment of gut colonization. * ff= force feed, p.ff= post force feed

To confirm the prolonged colonization stability of *K. pneumoniae* and *E. coli* strains in the larval gut, a group of larvae (n = 5×2= 10 per bacterial dose) was thrice force-fed with 10^6^ CFU/larvae and incubated until 120 h post last force-feed, i.e., 168 h after the first dose of bacteria. After 120 h incubation, the same protocol for CFU counting was performed as described above.

### 2.6 *K. pneumoniae* infection in the hemolymph of larvae

Larvae were divided into five groups (n = 5×2 = 10 per dose), with each group receiving log_10_-fold increasing bacterial dosages of *K. pneumoniae* 14520, i.e., 102 CFU/larvae to 10^6^ CFU/larvae. From the early exponential phase, bacterial suspension was centrifuged, washed twice with PBS, and adjusted to 10^8^ CFU/mL in PBS corresponding to 10^6^ CFU/larvae. Bacterial doses were injected through the middle proleg into the larvae using a 10 µL ALS syringe, and the larvae were incubated at 37°C. The survival of the larvae was observed every 24 h, and Kaplan-Meier curves were generated.

### 2.7 Selection of bacteriophages dosage for *K. pneumoniae* strains

To evaluate the effect of bacteriophages on *K. pneumoniae* strains, growth curves were generated. A 100 µL bacterial suspension of 0.5 McFarland was added in each well of 96-well plate. The bacteriophages UZG4 and UZG13 (Table 2), with titers of 10^3^ PFU/well (data not shown as it was not effective), 10^5^ PFU/well and 10^7^ PFU/well individually and in the form of a cocktail, were added per well of 96 well plates containing bacterial suspensions of *K. pneumoniae* strains (Table 1). The optical density at 600 nm (OD_600_) was measured every 10 min for 22 hours using the plate reader Infinite 200 Pro (Tecan Group AG, Männerdorf, Switzerland) with an ongoing shaking incubation at 37°C.

### 2.8 Decolonization of the larvae gut

Each group of larvae (n = 3×2 = 6) was force-fed thrice with *K. pneumoniae* strains Kp 14520, Kp 419614 and ATCC 700603, with a dose of 10^6^ CFU/larvae (Figure 2). Post 24 h of last force-fed, each group was one time force-fed with either a phage cocktail of UZG4 and UZG13 (10^7^ PFU/larvae), 4 mg/L of ciprofloxacin or 2 mg/L meropenem (inhibiting the growth of all *K. pneumoniae* strains in this study) to compare the effectiveness of the phage cocktail to the antibiotic treatment. The phage cocktail with high titer was used due to the one-time force feeding and its ability to hinder the growth in all *K. pneumoniae* strains (Figure S2). After 24h of incubation time, larvae were anesthetized on ice for 30 min and their guts were collected. The gut was crushed in 1 x PBS, 1:10 serially diluted, and plated on MacConkey agar for CFU counting.

**Figure 2.**
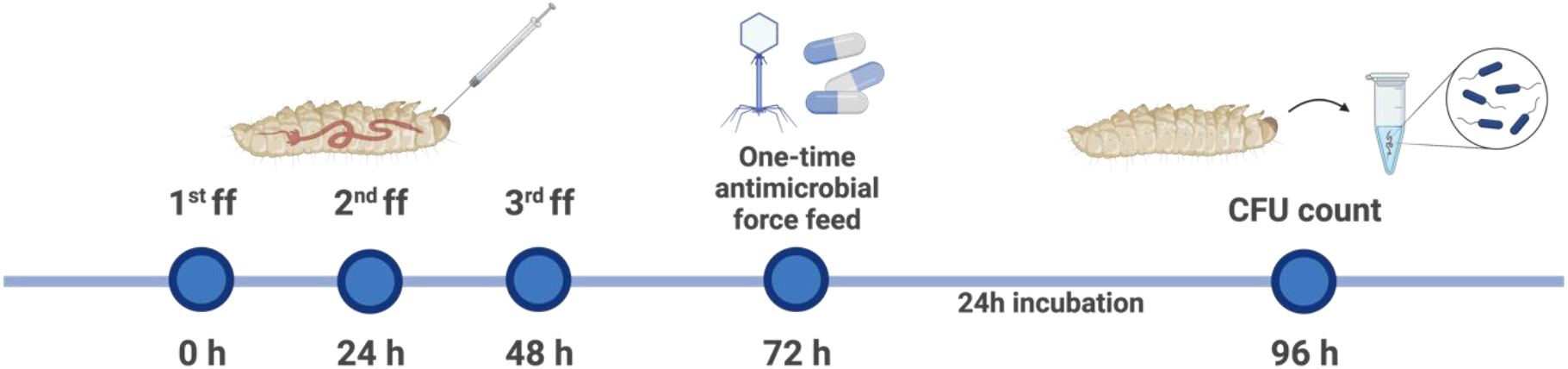
Timeline depicting the methodology adopted for the decolonization of colonized gut bacteria using antimicrobial treatment. * ff= force feed, p.ff= post force feed

### 2.9 Transmission electron microscopy

The bacteriophages on the bacterial cell were visualized using transmission electron microscopy (TEM) (Figure S3). A 0.5 McFarland bacterial suspension of ATCC 700603 was mixed with 10^7^ PFU/mL bacteriophage cocktail of UZG4 and UZG13. The suspension was incubated at 37°C at 160 RPM shaking for 4 h. After the incubation, the suspension was centrifuged at 10000 RCF for 1 hour. The pellet was fixed using 2.5% glutaraldehyde (SERVA Electrophoresis GmbH, Heidelberg, Germany) mixed with 0.1 M cacodylate buffer (SERVA Electrophoresis GmbH, Heidelberg, Germany) for 1 h.

## 3 Statistical analysis

Growth curve data were processed with Microsoft Excel (Microsoft Corporation, Redmond, USA). All data were further processed and analyzed using GraphPad Prism 9 (GraphPad Software Inc., San Diego, USA).The Kaplan Meier curves were compared using Log-rank (Mantel-Cox) test. The gut colonization CFU count was analyzed using the Kruskal-Wallis test. The gut decolonization CFU count was analyzed using the Mann-Whitney test. Two-sided confidence intervals of 5-95% were assumed. *P*-values of <0.05 were considered significant.

## 4 Results

### 4.1 Survival and colonization stability of larvae

Larvae were fed with increasing concentrations of bacteria to determine the optimal dose for colonization in the larvae gut without causing high mortality. The Kp 14520 isolate was force-fed thrice to the larvae at different doses, and ≥ 80 % of the larvae survived the colonization even in the highest dose group. The survival rate was 97 % with the force-feeding dose of 10^2^ CFU/larvae and 80 % with the bacterial dose of 10^6^ CFU/larvae group at 72 h (Figure 3a). The bacterial doses of 10^5^ CFU/larvae and 10^6^ CFU/larvae showed significant difference in survival compared to control (MH media) and 10^2^ CFU/larvae bacterial dosage (Table S1).

**Figure 3.**
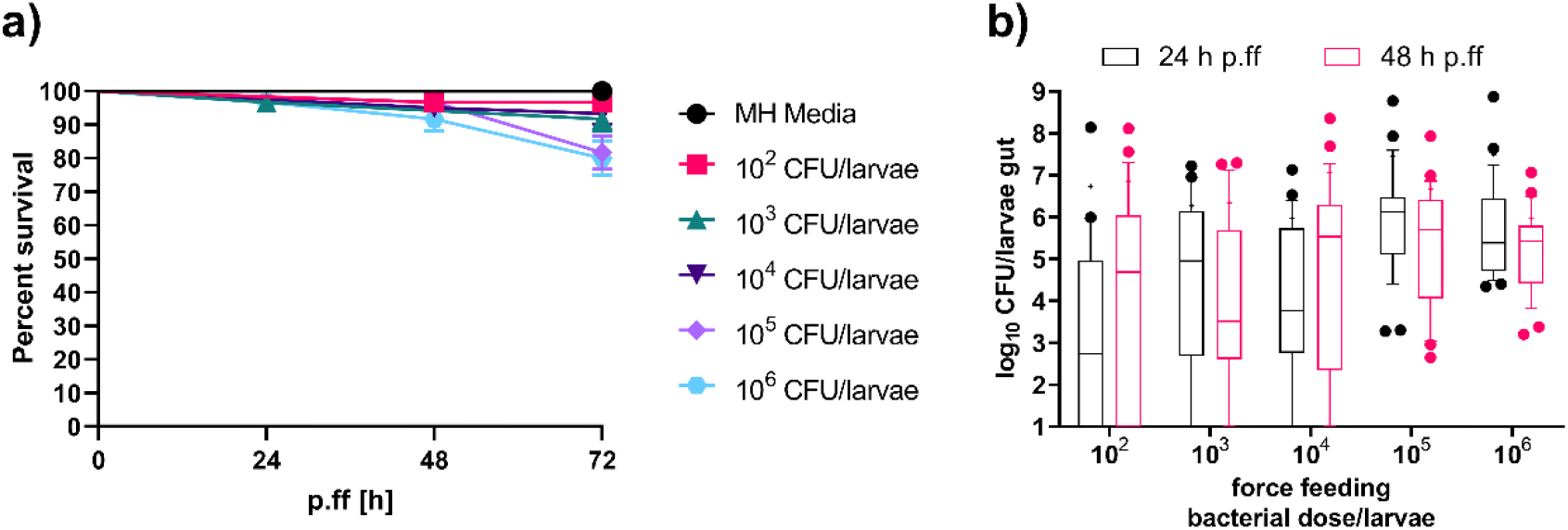
Survival curve of the larvae force-fed (a) and CFU count of the colonized bacteria post 24 h and 48 h of last force-feeding (b) with increasing bacterial doses of Kp 14520. (a) Larvae were force-fed at 0 h, 24 h, and 48 h with the bacterial doses indicated in the legend. The survival of the larvae was noted at 24 h, 48 h, and 72 h post force-fed. Each experiment was repeated thrice with 20 larvae per group (n = 60). (b) Box and whiskers graph of the gut colonization of the different larvae groups post 24 h and 48 h of last force-feeding. The Gram-negatives were selected on MacConkey agar. The experiment was performed thrice, with six larvae per group on the first attempt and ten larvae per group on the second and third attempts (n = 26). The boxes represent the 90-10 percentile, the line represents the median, + represents the mean and dots represent values outside the percentile.

The stability of colonization was assessed by determining the number of CFU in the gut of alive larvae from all dosage groups post 24 and 48 h last force-feeding. All larvae groups showed colonization regardless of the bacterial dose concentration force-fed, however with varying levels of variability (Figure 3b). At both time points, the fed bacterial dosages of 10^5^ and 10^6^ CFU/larvae showed the lowest variability in CFU count in the gut, indicating more stable colonization compared to lower dosages. The median CFU numbers of 10^5^ and 10^6^ CFU/gut were approximately similar to the amount of bacteria fed at each dosage. The bacterial doses of 10^5^ CFU/larvae and 10^6^ CFU/larvae showed significant difference in terms of CFU count compared to 10^2^ CFU/larvae (Table S1). Furthermore, 10^5^ CFU/larvae exhibited significant difference in CFU count compared to 10^3^ CFU/larvae and 10^4^ CFU/larvae (Table S1). Consequently, further colonization experiments proceeded with the colonization dosage of 10^5^ and 10^6^ CFU/larvae.

To further differentiate between colonization and infection, the larvae were injected with different doses of Kp 14520 into the hemolymph to mimic a possible escape of the bacteria from the gut. Results showed that 10^6^ CFU/larvae and 10^5^ CFU/larvae were sufficient to kill all larvae within 24 h of inoculation (Figure 4). The survival rate of 58% and 25% was observed with the infection dosage of 10^3^ and 10^4^ CFU/larvae, respectively. The minimum infection dose of 10^2^ CFU/larvae led to a survival rate of 80 %, similar to three times force-feeding with 10^6^ CFU/larvae (Figure 3a). Significant differences in survival were found in the 10^3^, 10^4^, 10^5^, and 10^6^ CFU/larvae injected groups in comparison with the PBS injected group (Table S1).

**Figure 4.**
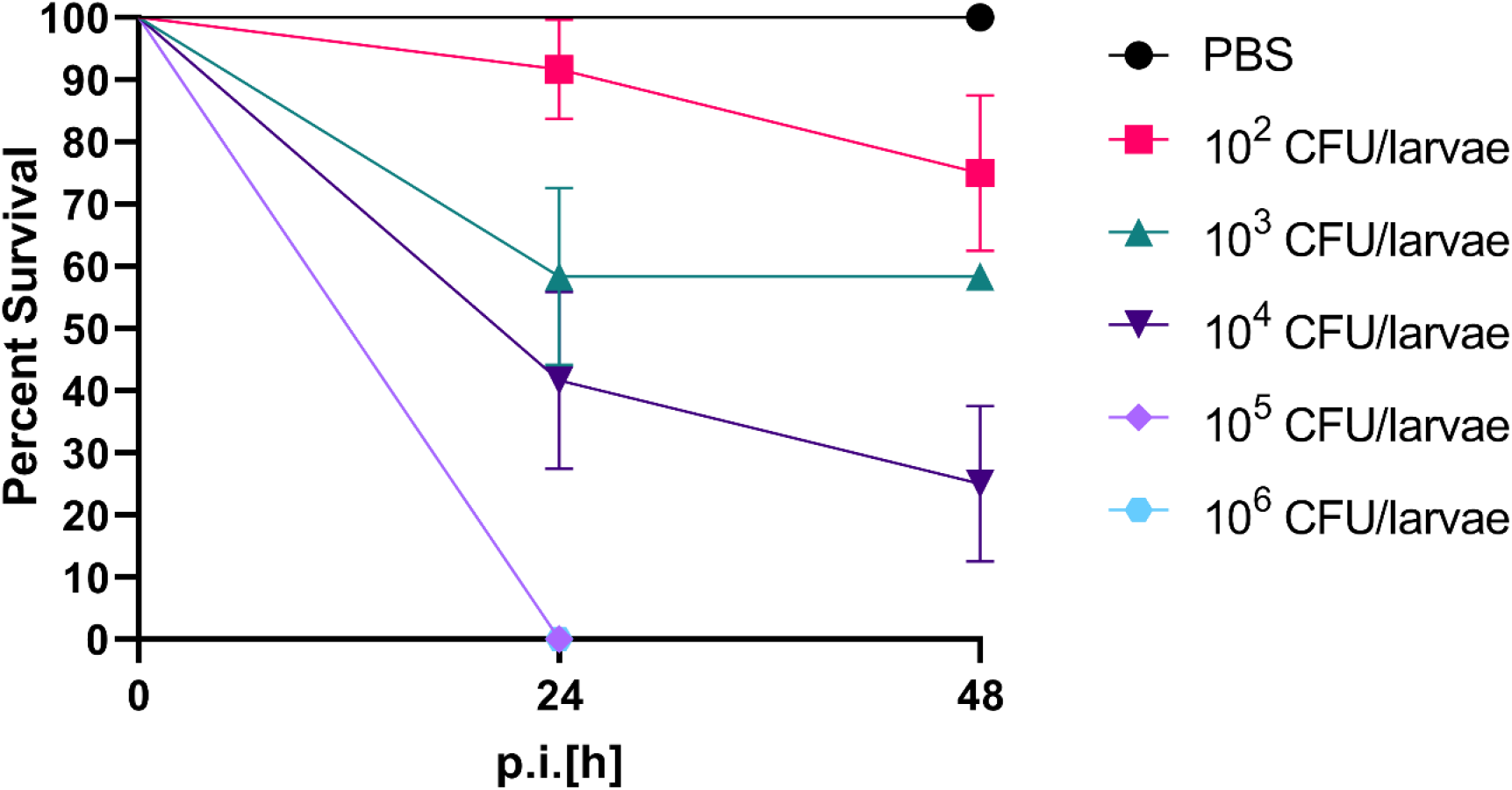
The Kaplan Meier curve of the larvae groups injected with bacterial doses in hemolymph through proleg. The 5 groups of larvae were injected with 10^2^, 10^3^,10^4^, 10^5,^ and 10^6^ CFU/larvae. The experiment was repeated twice with 6 larvae per group (n = 12). The error bars represent standard error (SE).

The effect of isolate and bacterial species variation on the stability of colonization was evaluated by colonizing larvae with either thrice 10^5^ or 10^6^ CFU/larvae of Kp 419614, ATCC 700603, ATCC 35218, Ec 208873 and Ec 280624. The survival of larvae force-fed with *K. pneumoniae* strains was 80%, except for the larvae group force-fed with ATCC 700603, which received a bacterial dose of 10^5^ CFU/larvae and had a survival rate of 70% (Figure 5a). A significant difference in survival was found between ATCC 700603 10^5^ CFU/larvae and MH media force-fed larvae (Table S1). In comparison to the *K. pneumoniae* strains, the larvae groups fed with *E. coli* strains showed greater mortality, with 50% and 60% survival recorded for ATCC 35218 and 280624, respectively (Figure 5b). The significant differences in survival of larvae force-fed 10^5^ CFU/larvae and 10^6^ CFU/larvae of *E. coli* strains compared to the MH medium were observed statistically as well (Table S1). CFU counts were determined 48 h post last force-feeding for all larval groups and showed colonization. The median colonization of the larvae with *K. pneumoniae* strains was 5 log_10_ CFU per larvae with bacterial dose of 10^6^ CFU/larvae (Figure 5c). On the other hand, larvae colonized with *E. coli* strains had lower median CFU counts (Figure 5d). When larvae were force-fed with 106 CFU/larvae of *E. coli* strains, the median CFU count was 4 log_10_ CFU/larvae, respectively. However, the mean CFU count of larvae that were force-fed with 10^6^ CFU/larvae of *E. coli* strains was the same as the amount force fed. Notably, some of the larvae were neither colonized with *K. pneumoniae* nor *E. coli*, regardless of the bacterial strain. The bacterial dose of 10^6^ CFU/larvae showed less variation compared to 10^5^ CFU/larvae so further experiments proceeded with 10^6^ CFU/larvae dosage.

**Figure 5.**
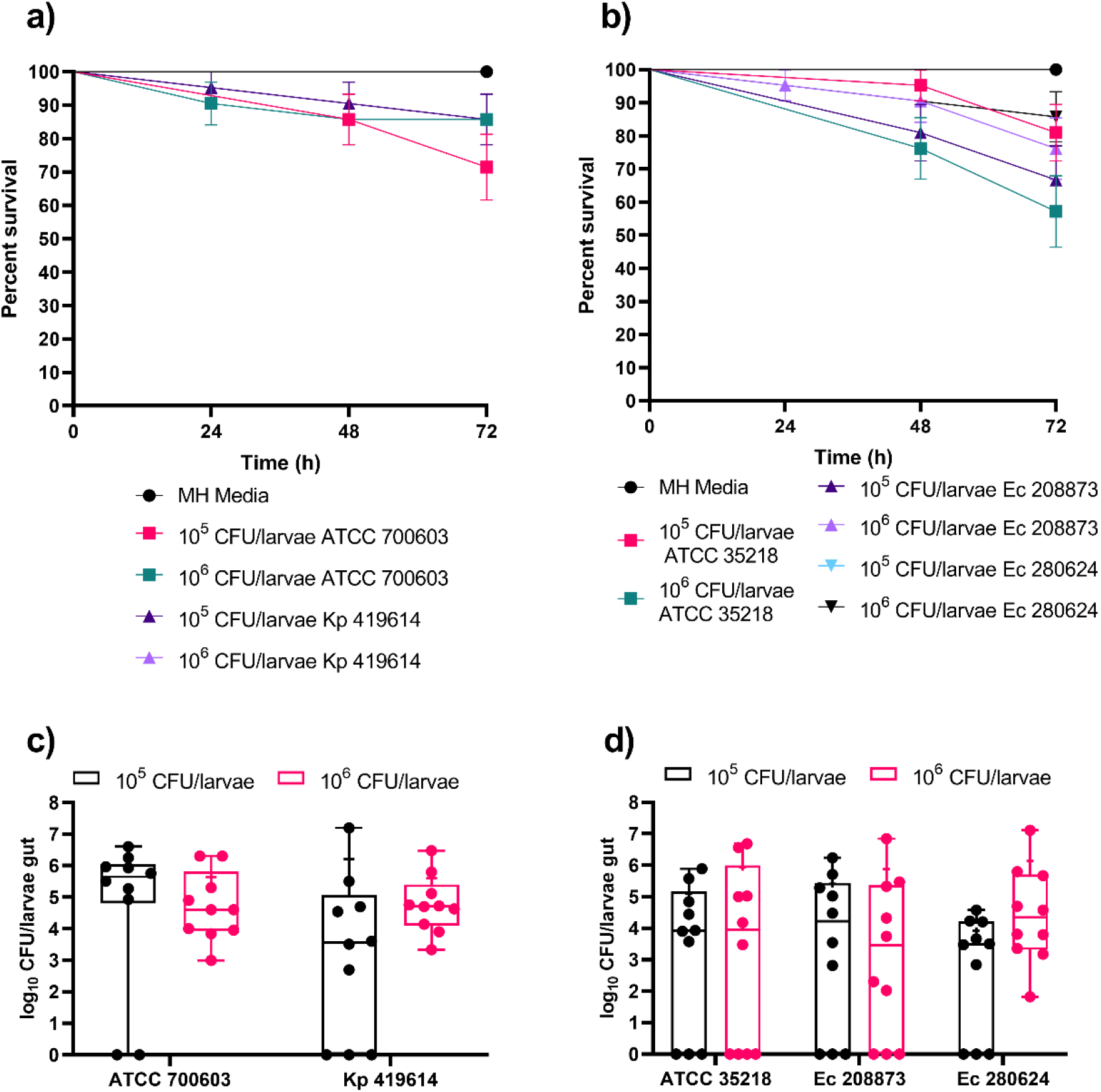
Survival curve of larvae colonized (a) and CFU count post 48h of last force-feed of the larvae gut colonized with *K. pneumoniae* ATCC 700603 and Kp 419614 (c). Survival curve of larvae colonized (b) and CFU count post 48h of last force-feed of the larvae gut colonized with *E. coli* ATCC 35218, isolates Ec 208873 and Ec 280624 (d). Larvae were colonized with 10^5^ and 10^6^ CFU/larvae bacterial doses of *K. pneumoniae* and *E. coli* strains (a, b). Three-force feedings were done and every 24 h survival was observed. The experiment was repeated thrice (6×2+1×9 = 21). The error bars represent standard error (SE). Larvae were divided into groups based on the bacterial dose force-fed to the larvae (c, d). Post 48 h of last force-feeding, larvae gut was isolated. The MacConkey agar was used for Gram-negative bacteria selection. The experiment was performed twice, with five larvae per group (n=10). The boxes represent 75 percentile, whiskers represent minimum to maximum, line represents median, + represents the mean and dots represent all data points.

The long-term stability of colonization and survival of larvae was assessed by force feeding with a thrice bacterial dosage of 10^6^ CFU/larvae of *K. pneumoniae* and *E. coli* strains each and incubating for 120 hours post the last force feed. The results showed a decrease in survival of 10-20% compared to the post-48 h colonization (Figure 5). For example, the survival of the larvae group colonized with ATCC 700603, Kp 419614, ATCC 35218 and Ec 280624 was 67%, 61%, 72%, and 72%, respectively (Figure 6a). However, the larvae colonized with Ec 208873 showed the least survival, which was only 39% post 120 hours of last force-feed. The survival of the larvae force-fed with *K. pneumoniae* and *E. coli* strains was significantly different from the survival of larvae force-fed MH media (Table S1). Despite the lower survival rate, the alive larvae still exhibited colonization, with the median CFU count of the larvae colonized with *K. pneumoniae* ranging from approximately 6 log_10_ to 7 log_10_, while the *E. coli* strains were approximately 5 log_10_ (Figure 6b). These results indicate that the colonization remained stable in surviving larvae even after the incubation of 120 hours post force feed in the larvae gut. The *K. pneumoniae* strains showed less variation compared to the *E. coli* strains so further decolonization experiment was only performed with *K. pneumoniae* strains.

**Figure 6.**
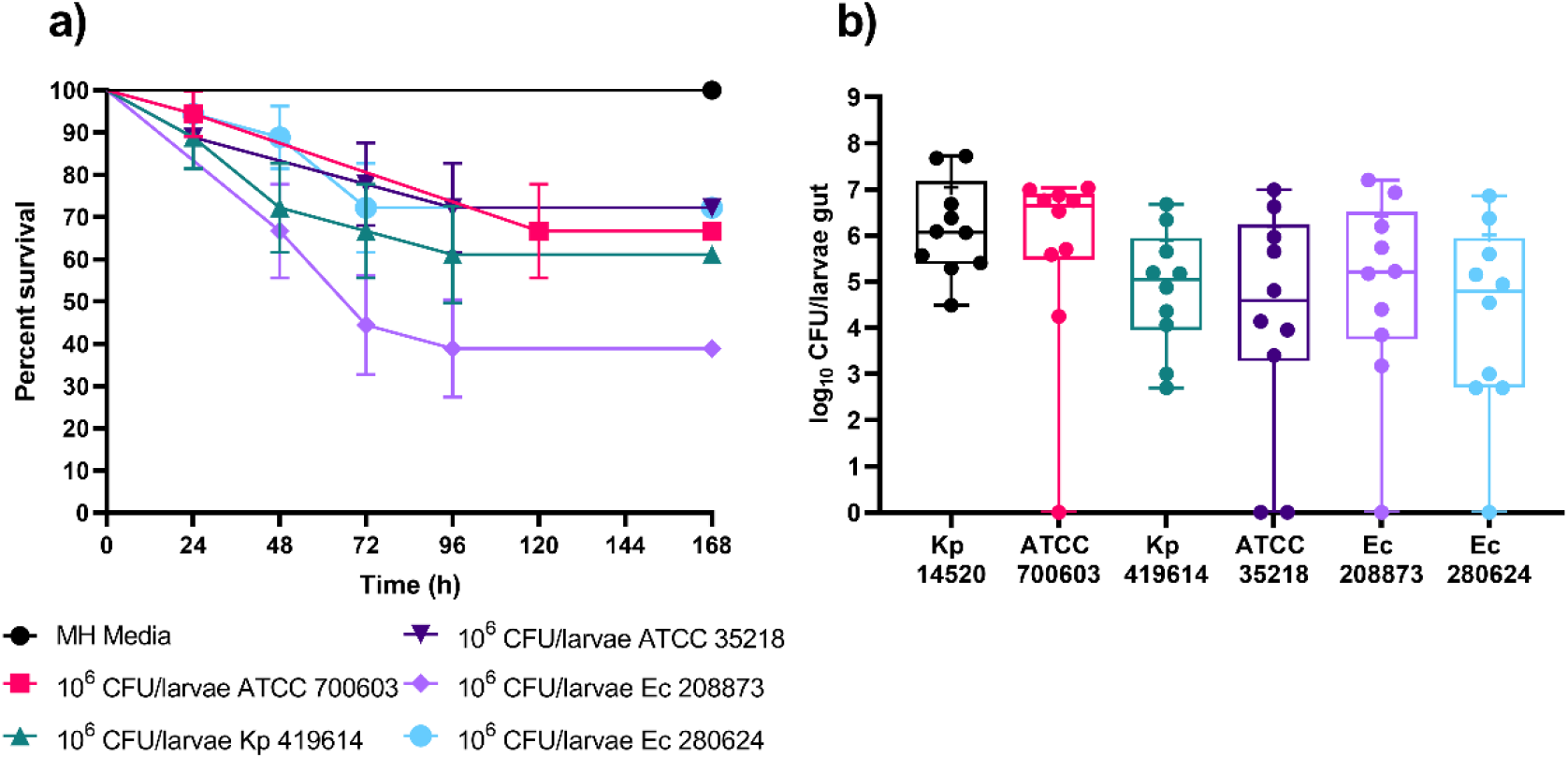
Survival curve of larvae colonized (a) and CFU count post 120 h of last force-feed (b) of the larvae gut colonized with *K. pneumoniae* and *E. coli* strains. Larvae were colonized with 10^6^ CFU/larvae bacterial dose. Larvae were force fed thrice and monitored for survival every 24 h. The experiment was repeated thrice (6×3= 18). The error bars represent standard error (SE). Larvae were divided into different groups based on the bacterial strain force-fed to the larvae (b). Post 120 h of last force-feeding, larvae gut was isolated and plated for CFU counting on the MacConkey agar. The experiment was performed twice, with five larvae per group (n=10). The boxes represent 75 percentile, whiskers represent the minimum and maximum, line represents the median, + represents the mean and dots represent all data points.

### 4.2 Decolonization of larvae gut

The bacteriophage susceptibility toward the selected bacterial strains was tested using growth curves (Figure S2). The individual bacteriophages UZG4 and UZG13 showed high efficacy in inhibiting selective strains only such as ATCC 700603 and Kp 419614, respectively. Whereas, the UZG4 and UZG13 bacteriophage cocktail of 10^7^ PFU/larvae hindered the bacterial growth (*in vitro*) in all strains such as ATCC 700603, Kp 14520 and Kp 419614 (Figure S2). For *in vivo* decolonization, larvae were thrice force-fed at 24 h intervals with a bacterial dose of 10^6^ CFU/larvae. Post 24 h of the last force feed, the first group of larvae were one-time fed with UZG4 and UZG13 phage cocktail of 10^7^ PFU/larvae, the second group was one-time fed with 4 mg/L ciprofloxacin, and the third group was one-time force-fed with 2 mg/L meropenem, in 10 µL of MH media (Figure 7). Results indicated that the bacteriophage cocktail significantly reduced median CFU counts compared to control such as 3 log_10_, 4 log_10_, and 2 log_10_ of *K. pneumoniae* strains ATCC 700603, 419614, and 14520, respectively (Table S1). However, one-time force feeding of antibiotics also showed a reduction in median CFU count but no significant reduction compared to control was found except 1 log_10_ reduction in group treated with ciprofloxacin in ATCC 700603 (Table S1). Moreover, significant difference between the phage therapy and ciprofloxacin treatment was observed in the larvae colonized with ATCC 700603 and Kp 49614 (Table S1). Similarly, significant difference between phage therapy and meropenem treatment was observed in the larvae force fed with *K. pneumoniae* strains 14520 and 419614 (Table S1). One-time application of the bacteriophage cocktail reduced CPE colonization. Mock treatment using the same concentrations of phages and antibiotics revealed that all larvae survived.

**Figure 7.**
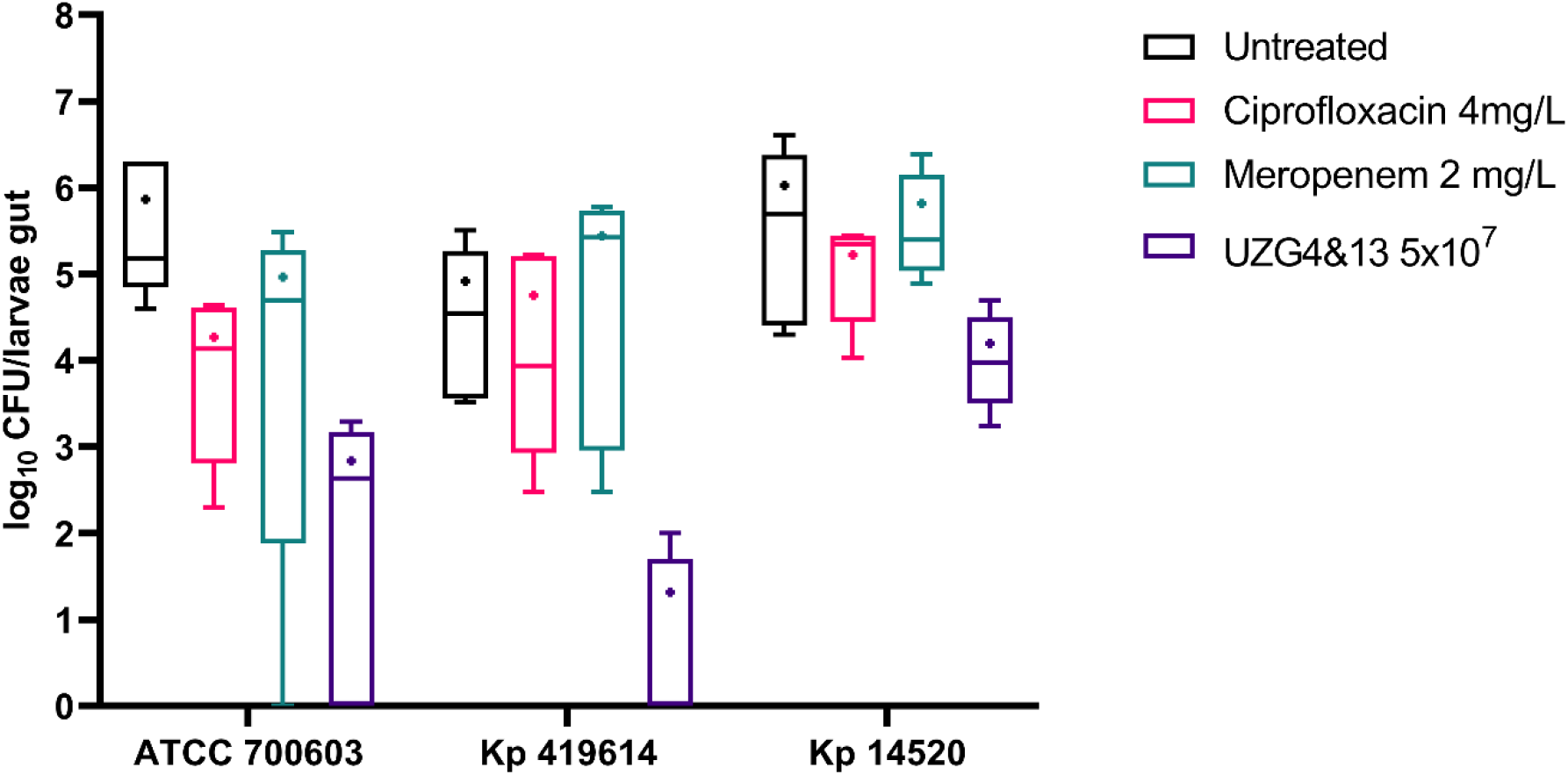
Decolonisation of the *K. pneumoniae* strains from the larvae gut using bacteriophage cocktail, ciprofloxacin and meropenem. The larvae were colonized with *K. pneumoniae* strains. After the colonization, larvae were one time force-fed with 10^7^ PFU/larvae bacteriophage cocktail, 4 mg/L ciprofloxacin and 2 mg/L meropenem and incubated for 24 h. CFU counts were evaluated after incubation. The experiment was repeated twice with 3 larvae per group (n=6). The boxes represent the 90 to 10 percentile, the line represents the median and + represents the mean.

## 5 Discussion and conclusion

This study investigated for the first time the use of *Galleria mellonella* larvae as a model for gut colonization of *K. pneumoniae* and *E. coli*. In addition, a one-time bacteriophage cocktail was administered through force-feeding to reduce *K. pneumoniae* colonization in larvae gut. The results showed that *K. pneumoniae* and *E. coli* were able to colonize the larvae gut for 120 hours after force-feeding in the majority of the larval population and with different bacterial strains. A bacteriophage cocktail was able to significantly reduce the colonization of *K. pneumoniae* in the gut compared to antibiotic treatment.

Our results are in contrast to previous research, in which oral force-feeding of Gm larvae with Shigella species was unsuccessful in establishing permanent colonization, with the larvae rapidly dying after oral bacterial administration (Barnoy et al., 2017a). The varying levels of pathogenicity demonstrated by various bacteria in different organisms may be the cause (Barnoy et al., 2017b). This can be attributed to the differences in the host’s immune system, which can lead to varying levels of susceptibility to a particular pathogen. Furthermore, the differences in the genetic makeup of the bacteria may also be responsible for the different levels of pathogenicity, as some bacteria may have evolved to be more harmful to certain hosts. The strain variation plays a vital role in distinct virulence between different isolates of the same bacterial species. The microbiome of *Galleria mellonella* is dominated by Enterococcus species (Allonsius et al., 2019) and *K. pneumoniae* and *E. coli* are not part of the native larval microbiome. The larvae are therefore more tolerant to high doses of Gram-positive bacteria compared to Gram-negative bacteria. In this study, a dose of 10^5^ CFU/larvae of Kp 14520 was sufficient to kill the entire larval population within 24 hours of infection in the hemolymph. Additionally, injection of 10^2^ CFU/larvae bacterial dose in hemolymph was enough to reduce the larval population to less than 80% after 48 hours of incubation. However, force-feeding of the high bacterial dose of 10^6^ CFU/larvae did not reduce the larvae population to less than 50 % post 120 h incubation post last force-feed. The only exception was Ec isolate 208873, which may have been due to strain variation. Overall, colonization of the larvae gut with all bacterial strains was successful and stable over time in alive larvae. However, some of the larval population showed mortality over time maybe because of bacteria overcoming the immune system and causing infection.

The emergence of antibiotic-resistant pathogens has increased the urgency of finding alternative treatments for bacterial infections. Bacteriophages, viruses that specifically target bacterial species, have become an attractive option due to their specificity and lack of toxicity to other cells. However, the use of bacteriophages for intravenous treatment is challenging due to the release of cell debris and endotoxins released upon bacterial lysis. Fortunately, the gut has a greater tolerance to endotoxins compared to the sterile parts of the body, making the gut an ideal environment for treatment with bacteriophages. In this study, bacteriophages exhibited higher efficacy in terms of growth inhibition against CPE. Our study has indicated that phages have a remarkable ability to inhibit bacterial growth, performing more consistently than antibiotics, particularly in CPE decolonization.

Recent decolonization studies have used mice models to investigate the efficacy of bacteriophage therapy for decolonizing intestinal *K. pneumoniae*. Mice were pre-treated with meropenem (Fang et al., 2022) or germ-free mice were used (Liu et al., 2022) and then infected with *K. pneumoniae* to simulate a colonization state. Upon development of colonization, the mice were subsequently treated with different concentrations of phages and monitored for decontamination efficacy. Results showed that the phage treatment resulted in significantly decreased levels of *K. pneumoniae* in the gut microbiota compared with both the untreated groups. In our study, the one-time phage cocktail was able to significantly reduce the colonization of *K. pneumoniae* strains within the gut, in comparison to the effect of standard antibiotics, while simultaneously not killing the larvae. Interestingly, in this regard, the administration of antibiotics was observed to have less to no significant reduction in comparison to the bacteriophages, despite the colonization with the non-resistant ATCC 700603. This could stem from the possibility of the bacteria forming biofilm. A biofilm mode of life adopted by many bacterial species, such as *K. pneumoniae*, enables the bacteria to attach to the surface of the gut (de Vos, 2015) thus making them highly tolerant to antibiotics.

The gut colonization of mice has been widely studied in recent years, yet the results have been mixed. While some studies have used low-complexity-microbiota mice (Göttig et al., 2015) or antibiotics treatment such as ampicillin to establish colonization (Xenofontos et al., 2022), direct feeding to achieve colonization has proven to be challenging due to the complexity of the mice microbiome and immune system. As such, the high usage of mice in number of experiments without certainty of successful colonization is an issue of ethical concern. We presented a novel protocol to achieve gut colonization of Gm larvae using three force-feedings, without the need for any antibiotics pre-treatment or germ-free gut. This protocol can be further tested with limited numbers of mice, taking advantage of the ease of use, quick processing, and high number of replicates offered by the larvae model. While the larvae model cannot completely replace the mammalian model, its advantages of ease of use, quick processing and high number of replicates can help to reduce the unnecessary consumption of mice and provide an effective alternative for research purposes.

*Galleria mellonella* is an effective alternative model for gut colonization and decolonization studies. It is also shown that bacteriophage treatments can be utilized to decontaminate the larval gut. However, this study is limited to the observation of *K. pneumoniae* and *E. coli* strains with regard to gut colonization and *K. pneumoniae* strains related to phage-mediated decolonization. To develop a more comprehensive understanding, future studies should incorporate more bacterial species. Additionally, exploration of phages that are more effective for bacterial decontamination should be explored. Furthermore, experiments should be conducted to determine the effect of sequential administrations of phages with and without antibiotics, as well as to monitor bacterial and phage resistance patterns after treatments. Overall, more research needs to be done to enhance the applicability of this model for gut colonization and decontamination using novel antimicrobials, to successfully decontaminate resistant pathogens from the gut.

## 6 Acknowledgments

## 8 Declaration

