## Supplementry data for "Bacteriophage-mediated decolonization of *Enterobacteriaceae* in a novel *Galleria mellonella* gut colonization model"

### 1 Supplementary Data

**Table S1** Statistical analysis and significantly different groups of the study.

| **Target groups** | **Significance**<br>$p<0.05$ (*), $p<0.01$ (**), $p<0.001$ (***), $p<0.0001$(****) |
|---|---|
| **Figure 3a: Statistical analysis using Log-rank (Mantel-Cox) test** | |
| MH media vs $10^5$ CFU/larvae | * |
| MH media vs $10^6$ CFU/larvae | ** |
| $10^2$ CFU/larvae vs $10^5$ CFU/larvae | * |
| $10^2$ CFU/larvae vs $10^6$ CFU/larvae | ** |
| $10^4$ CFU/larvae vs $10^6$ CFU/larvae | * |
| **Figure 3b: Statistical analysis using the Kruskal-Wallis test** | |
| $10^2$ CFU/larvae (24h) vs $10^5$ CFU/larvae (24h) | **** |
| $10^2$ CFU/larvae (24h) vs $10^6$ CFU/larvae (24h) | ** |
| $10^2$ CFU/larvae (24h) vs $10^5$ CFU/larvae (48h) | * |
| $10^4$ CFU/larvae (24h) vs $10^5$ CFU/larvae (24h) | * |
| $10^3$ CFU/larvae (48h) vs $10^5$ CFU/larvae (24h) | * |
| **Figure 4: Statistical analysis using Log-rank (mantel-Cox) test** | |
| PBS vs $10^3$ CFU/larvae | * |
| PBS vs $10^4$ CFU/larvae | *** |
| PBS vs $10^5$ CFU/larvae | **** |
| PBS vs $10^6$ CFU/larvae | **** |
| $10^2$ CFU/larvae vs $10^4$ CFU/larvae | ** |
| $10^2$ CFU/larvae vs $10^5$ CFU/larvae | **** |
| $10^2$ CFU/larvae vs $10^6$ CFU/larvae | **** |
| $10^3$ CFU/larvae vs $10^5$ CFU/larvae | ** |
| $10^3$ CFU/larvae vs $10^6$ CFU/larvae | ** |
| $10^4$ CFU/larvae vs $10^5$ CFU/larvae | * |
| $10^4$ CFU/larvae vs $10^6$ CFU/larvae | * |
| **Figure 5a: Statistical analysis using Log-rank (mantel-Cox) test** | |

| | |
|---|---|
| MH media vs $10^5$ CFU/larvae ATCC 700603 | ** |
| **Figure 5b: Statistical analysis using Log-rank (mantel-Cox) test** | |
| MH media vs $10^5$ CFU/larvae ATCC 35218 | * |
| MH media vs $10^6$ CFU/larvae ATCC 35218 | *** |
| MH media vs $10^5$ CFU/larvae Ec 208873 | ** |
| MH media vs $10^6$ CFU/larvae Ec 208873 | * |
| MH media vs $10^5$ CFU/larvae Ec 280624 | * |
| **Figure 6a: Statistical analysis using Log-rank (mantel-Cox) test** | |
| MH media vs $10^6$ CFU/larvae ATCC 700603 | ** |
| MH media vs $10^6$ CFU/larvae Kp 419614 | ** |
| MH media vs $10^6$ CFU/larvae ATCC 35218 | * |
| MH media vs $10^6$ CFU/larvae Ec 208873 | **** |
| MH media vs $10^6$ CFU/larvae Ec 280624 | * |
| **Figure 7: Statistical analysis using the Mann-Whitney test** | |
| Positive control vs 4 mg/L ciprofloxacin (ATCC 700603) | ** |
| Positive control vs UZG 4&13 (ATCC 700603) | ** |
| Positive control vs UZG 4&13 (Kp 419614) | ** |
| Positive control vs UZG 4&13 (Kp 14520) | * |
| Ciprofloxacin 4 mg/L vs UZG 4&13 (ATCC 700603) | * |
| Ciprofloxacin 4 mg/L vs UZG 4&13 (Kp 419614) | * |
| Meropenem 2 mg/L vs UZG 4&13 (Kp 419614) | ** |
| Meropenem 2 mg/L vs UZG 4&13 (Kp 14520) | ** |

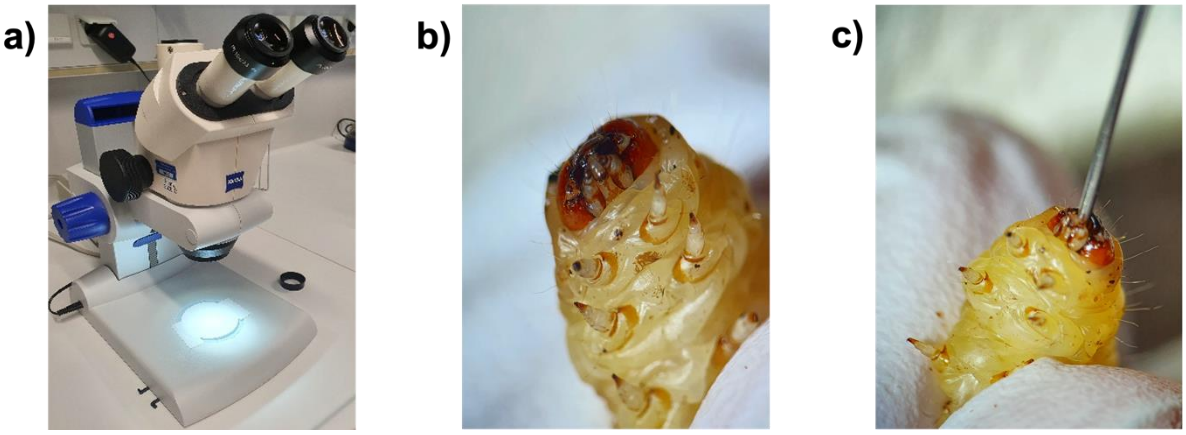


**Figure S1** (a) the Zeiss Stemi 2000-C zoom microscope. (b) the larvae's mouth under a microscope stereo-zoom microscope. (c) the larvae force-feeding under a microscope using a ALS blunt-end syringe.

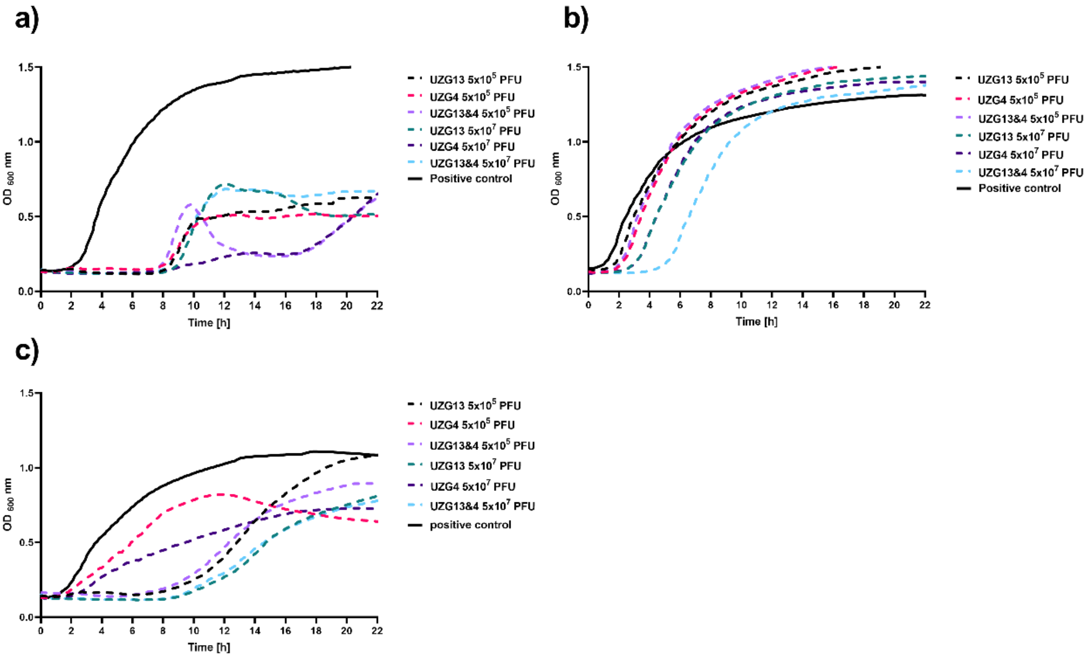


**Figure S2** The effect of different bacteriophages and titers on *K. pneumoniae* strains ATCC 700603 (a), 14520 (b) and 419614 (c) *in vitro*. The 100 µL bacterial suspensions of 0.5 McFarland were added into wells of 96 well microtiter plate. The 100 µL of $10^5$

and $10^7$ PFU/well of UZG4 and UZG13 bacteriophages individually and in form of cocktail were added into the well containing the bacterial suspension.

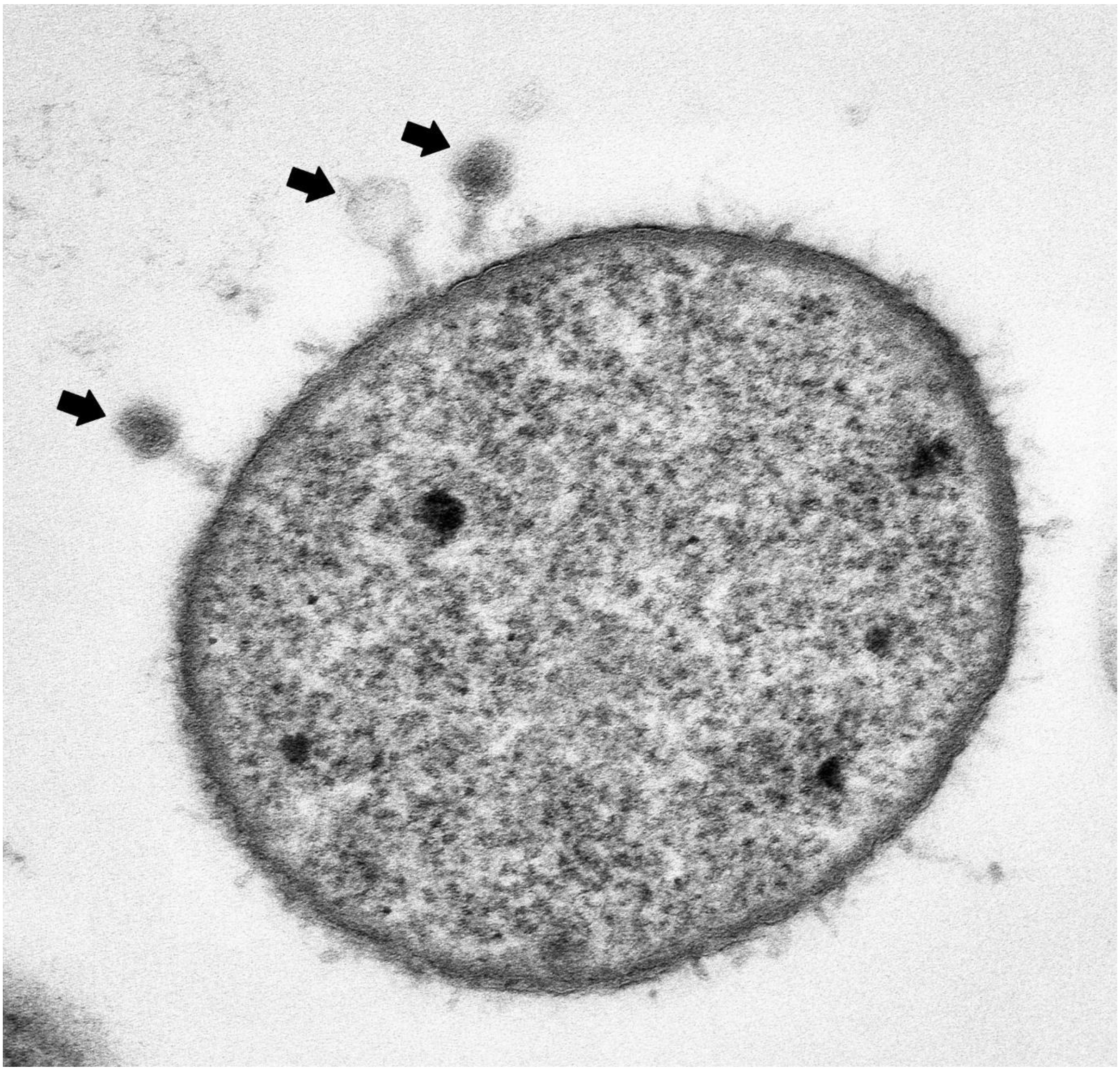

**Figure S3** Electron microscopic image of myoviridae bacteriophages UZG4 and UZG13 (arrows) attacking a *K. pneumoniae* ATCC 700603 bacterial cell. Magnification 85.000x, Scale bar: 200 nm
